# Estimating dispersal kernels using genetic parentage data

**DOI:** 10.1101/044305

**Authors:** Michael Bode, David Williamson, Hugo Harrison, Nick Outram, Geoffrey P. Jones

## Abstract

Dispersal kernels are the standard method for describing and predicting the relationship between dispersal strength and distance. Statistically-fitted dispersal kernels allow observations of a limited number of dispersal events to be extrapolated across a wider landscape, and form the basis of a wide range of theories and methods in ecology, evolution and conservation. Genetic parentage data are an increasingly common source of dispersal information, particularly for species where dispersal is difficult to observe directly. It is now routinely applied to coral reef fish, whose larvae disperse over many kilometers and are too small to follow directly. However, it is not straightforward to estimate dispersal kernels from parentage data, and existing methods each have substantial limitations. Here we develop and proof a new statistical estimator for fitting dispersal kernels to parentage data, applying it to simulated and empirical datasets of reef fish parentage. The method incorporates a series of factors omitted in previous methods: the partial sampling of adults and juveniles on sampled reefs; the existence of unassigned dispersers from unsampled reefs; and post-settlement processes (e.g., density dependent mortality) that follow dispersal but precede parentage sampling. Power analyses indicate that the highest levels of sampling currently used for reef fishes is sufficient to fit accurate dispersal kernels. Sampling is best distributed equally between adults and juveniles, and over more than ten populations. Importantly, we show that accounting for unsampled or unassigned individuals – including adult individuals on partially-sampled and unsampled patches – is essential for a precise and unbiased estimate of dispersal.

## Introduction

The pattern and strength of dispersal play a defining role in ecology and evolution, particularly in ecosystems where suitable habitat is patchily distributed (Clobert *et al.* 2001; Bullock *et al.* 2002). However, dispersal is a difficult process to observe, particularly when dispersing individuals are small, numerous and hard to follow (Cowen & Sponaugle 2009; Jones *et al.* 2009). A wide variety of approaches have been developed to measure dispersal events. These include tracking individual trajectories using radio or GPS devices; mark-recapture across multiple patches; and inverse modelling using the location and species identity of propagules and adults (Clobert *et al.* 2001; Bullock *et al.* 2002; Clark *et al.* 2008; Broquet & Petit 2009). In recent years, population and individual-based genetic methods have begun to offer unique and powerful insights into dispersal (Jones *et al.* 2009; Broquet & Petit 2009).

Hypervariable molecular markers allow sampled juveniles to be assigned to sampled parents, and can thereby provide individual dispersal vectors. The approach was first used to estimate dispersal and gene flow in plant communities (Ellstrand & Marshall 1985), has been applied to birds (Woltmann *et al.* 2012), and mammals (Telfer *et al.* 2003). It is increasingly being used to observe larval dispersal events in marine fish metapopulations, particularly on coral reefs (Jones *et al.* 2005b; Almany *et al.* 2007; Planes *et al.* 2009; Christie *et al.* 2010; Buston *et al.* 2012; Harrison *et al.* 2012; Almany *et al.* 2013; D’Aloia *et al.* 2015). Parentage analysis has been used to conclusively answer open ecological questions (Almany *et al.* 2007; Jones *et al.* 2009), and provide suggestive evidence about the role of connectivity in spatial management (Harrison *et al.* 2012). However, parentage assignment data generally only report a small number of dispersal events, at a particular time, and between a subset of locations. Many questions in ecology and management tools in conservation demand a more expansive description of dispersal.

We therefore require statistically robust methods for extrapolating limited parentage data to predictions about dispersal in the broader landscape. The most common approach at present is to estimate how dispersal strength relates to the distance between habitat patches, since isolation-by-distance is a natural way to conceive of dispersal dynamics. Operationally, this involves fitting a parametric relationship – a dispersal kernel – to a set of observed dispersal events (Clobert *et al.* 2001; Nathan *et al.* 2012). Previous analyses have fit dispersal kernels to parentage datasets using a range of different methods (Jones *et al.* 2005a; Jones & Muller Landau 2008; Buston *et al.* 2012; Almany *et al.* 2013; Hopf *et al.* 2015; D’Aloia *et al.* 2015). At their most basic, these methods involve calculating the average distance travelled by juveniles that were sampled and assigned to parents (Woltmann *et al.* 2012), or fitting a regression line (Telfer *et al.* 2003; Buston *et al.* 2012) directly to the distances travelled. In essence, these methods use each parentage assignment as a datapoint relating distance to dispersal strength. While this assumption is broadly reasonable, the raw number of assignments should not simply be regressed against inter-patch distance. The fit of dispersal kernels to parentage data will be influenced by a set of factors that are generally not included in current fitting methods. (1) Even the most extensive parentage datasets contain only subsamples of the adult populations. The data therefore contains a large numbers of juveniles – generally the majority – that cannot be assigned to any parents. These unassigned juveniles could be the offspring of adults on “ghost patches”, that is, patches that are completely unsampled (Beerli 2004; Wang 2014). They could also be the offspring of unsampled adults from patches where adults were only partially sampled (Jones & Ardren 2003). Both options must be considered by the method. (2) Parentage datasets contain finite samples of the juvenile populations, and the datasets vary spatially in sampling intensity (Jones & Muller Landau 2008). (3) These juvenile samples were not collected immediately following the dispersal phase, and therefore are the result of dispersal, filtered by post-settlement mortality processes (Moran & Clark 2012). These factors – each the result of partial sampling – can have a dramatic effect on the interpretation of a given parentage sample. First, unassigned juveniles contain valuable information that should not be ignored. Second, partially-sampled and unsampled adult populations can substantially change estimates of kernel shape and mean dispersal distance (see Figure S1 for a simple illustration).

All previous estimates of larval dispersal kernels for coral reef fish have ignored at least one of these factors. In this paper, we propose a novel likelihood estimator for dispersal kernels that incorporates each, and demonstrate its application to a case study of reef fish larval dispersal on the Great Barrier Reef. We generate simulated genetic parentage datasets to investigate the power and statistical properties of our estimator under logistical constraints, which limit the number of juveniles sampled, the proportion of adults sampled at each patch, and the total number of patches sampled. Finally, we show that methods which do not incorporate these factors will produce biased and inaccurate estimates of dispersal.

## A likelihood function for parentage-based dispersal data

The metapopulation comprises *P* habitat patches, each with a population of *N*_*i*_ individuals (we assume an approximately equal sex ratio on all patches). The metapopulation consists of individuals in three stages. “Adults” are reproductively mature individuals that produce dispersers (e.g., larvae) that move according to a dispersal kernel. If these offspring arrive at a habitat patch, they become “settlers”, and attempt to recruit to the local population. At this point they suffer density-dependent mortality; those that survive become “juveniles” and may be sampled for parentage.

Juveniles are sampled from a subset **S**_**J**_ of patches, which has *s*_*J*_ elements. A proportion π_*i*_ of the adults on each reef *i* are sampled, with π_*i*_ = 0 for ghost patches. Each sampled juvenile is either assigned to a sampled adult, or classified as having unknown parentage. This count data populates matrix **M**, with the columns indicating the patch where the juvenile was sampled, and the row indicating the natal patch. The final row contains the number of unassigned juveniles. The count data in **M** are a sample from the juveniles on each patch, which are themselves samples of the settlers. We assume that both the juveniles and settlement pools are large, so that we can model recruitment and juvenile collection as samples with replacement. We also assume that each disperser, regardless of its origin, has an equal probability of successfully settling (e.g., there is no local adaptive advantage (Warner 1997) or cost to long duration/distance dispersal (Burgess *et al.* 2013a)).

The settlement pool on each patch is determined by a dispersal kernel ρ(*k*, *d*_*ij*_) which assumes dispersal is isotropic, spatially invariant, and based on the distance *d*_*ij*_ between patches *i* and *j*. The kernel is a probability density function whose shape is defined by the parameter set *k*. One commonly-used kernel is the generalised Gaussian function (Largier 2003; Bode *et al.* 2011; Nathan *et al.* 2012):

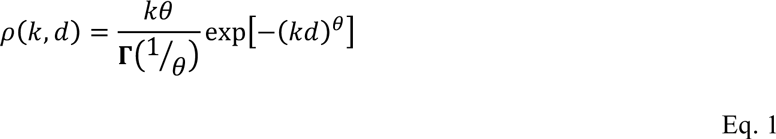

where *θ* is a shape parameter that yields the standard Gaussian when *θ* = 2, the Ribbens function when *θ* = 3, and the Laplacian or negative exponential when *θ* = 1. The coefficients that precede the exponential term in Eq. 1 normalise the function to ensure that it is a probability distribution. However, because these terms occur on both the numerator and denominator of terms in the likelihood function (Eq. 2), they do not affect the fit and could be safely ignored during fitting. Note that since the kernels are isotropic, the average displacement of an individual larvae is zero (since the displacement of every larvae that disperses northward is balanced by larva that travels south). We therefore define the mean dispersal distance as the expected distance travelled by a larvae in a single compass direction:

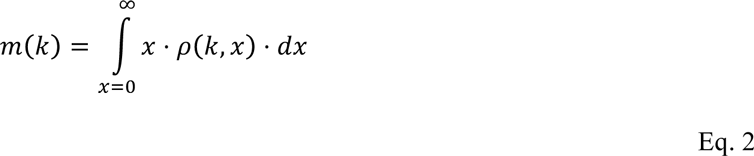

According to a particular dispersal kernel, the proportion of the settlers on patch *j* that come from sampled adults on reef *i* will be:

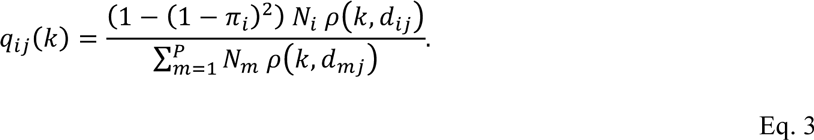

The numerator in Eq. 2 calculates the number of settlers who dispersed to patch *j*, that were created by at least one sampled parent on patch *i*. The denominator divides this by the total number of settlers arriving at patch patch *j*, from both sampled and unsampled adults, turning it into a probability that a sampled juvenile on *j* comes from patch *i*. This denominator recreates the effect of any density-dependent settlement mortality process that is neutral to the source of the settlers. Both the numerator and denominator could be modified by the per-capita fecundity of the females, the proportion of successfully fertilised eggs, and the mortality during the dispersal phase. However, if we assume that these do not vary between patches, they will not alter the fit. Equation 2 depends heavily on the population size on potential source patches, *N*_*j*_ (Fig 1a). These must therefore be either sampled, or estimated using observed densities on comparable sampled habitat.

**Figure 1:**
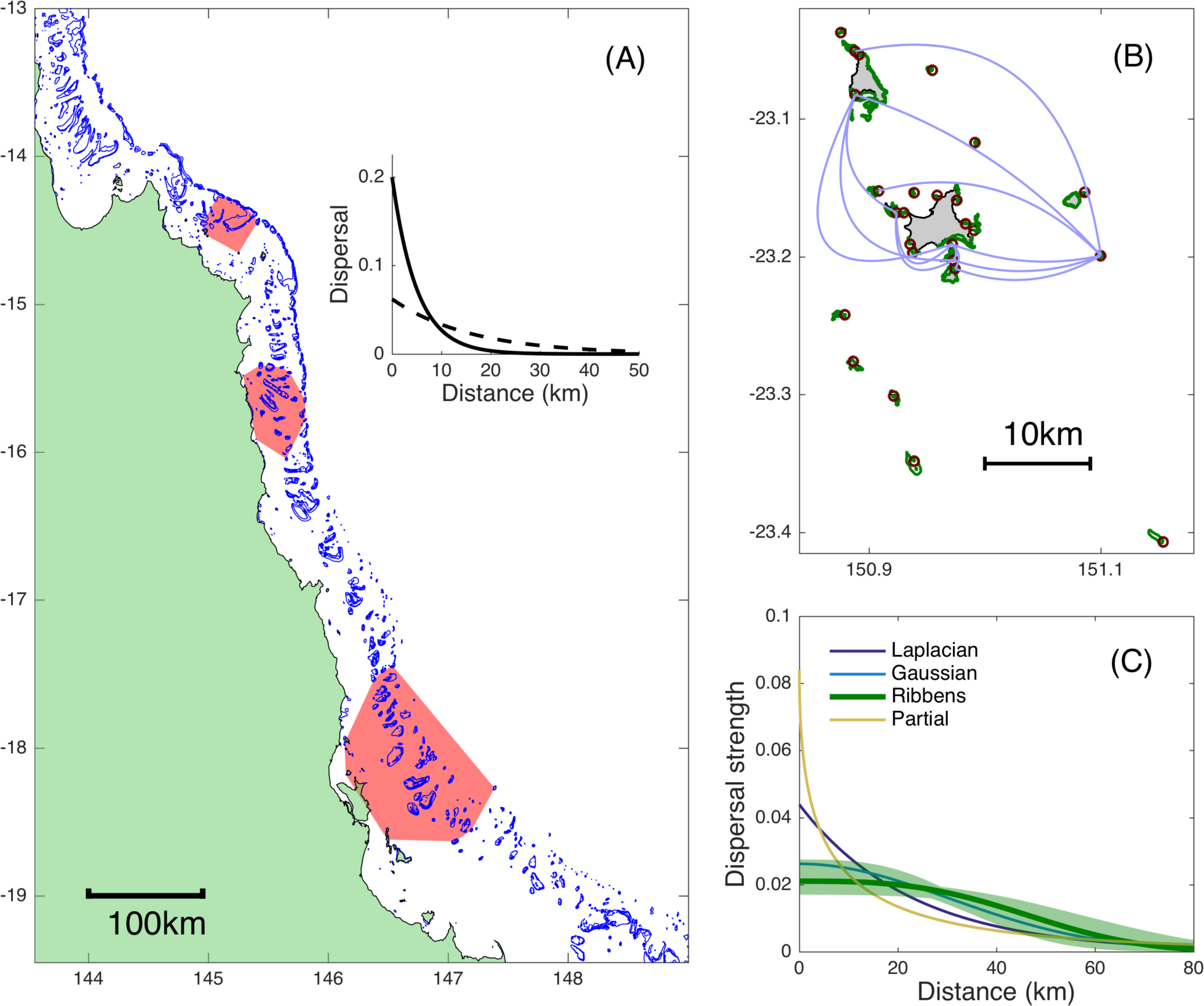
(A) Location of the reefs on the Great Barrier Reef, Australia, used for the power analysis simulations. Red polygons exemplify sets of 20, 40, and 60 reefs (from north to south) used to simulate parentage data and fit dispersal kernels. Inset panel shows the scale of the two dispersal kernels used for the power analyses. (B) Location of reefs in the Keppel Islands group, for the empirical fitting example. Green polygons indicate reefs; grey polygons are islands. Blue lines indicate larval exchange between reefs that were identified by parentage (directionality not shown). (C) Best-fit parameterisations for four larval dispersal kernel shapes. The Ribbens kernel (95% confidence intervals shaded in green) provided the best fit to the data.

In most parentage datasets, a large proportion of the juveniles cannot be assigned to any of the sampled adults. The dispersal kernel also predicts this proportion, which either come from ghost patches, or from unsampled adults on sampled patches:

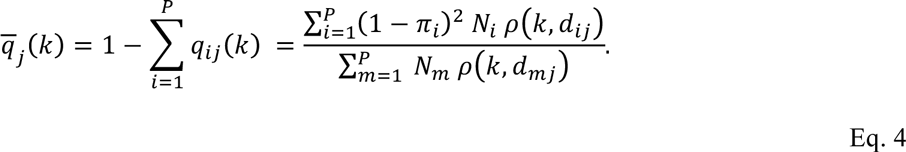

Applying Eq. 2 and Eq. 3 to the parentage matrix **M**, the log likelihood of observing the set of attributed and unattributed samples, given dispersal kernel *ρ*(*k*, *d*) is therefore:

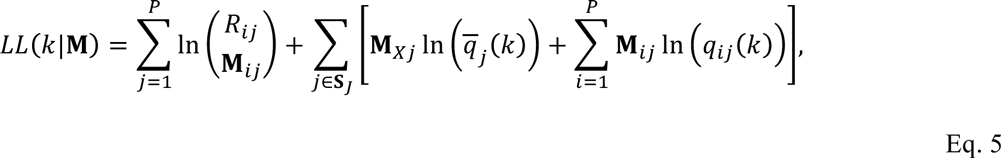

Where *R*_*ij*_ = Σ_*i*_ **M**_*ij*_, the total number of recruits sampled on patch *j*. The index variable *X* in **M**_*Xj*_ refers to the final row in the parentage matrix (i.e., unallocated juveniles, so *X* = *s*_*A*_ + 1). The very first term of Eq. 5, a sum over multinomial coefficients, does not change for different parameter sets, and should therefore be ignored since it can be large enough to pose numerical issues. Confidence intervals can be generated by repeatedly finding the maximum likelihood fit for bootstrap resamples of the data, using source reefs as resampling units.

## Applying the method to reef fish dispersal in the Keppel Islands Group

We begin by applying this estimator to an empirical genetic parentage dataset from the Keppel Islands Group, an archipelago of continental islands in the southern section of Australia’s Great Barrier Reef (Figure 1A). The genetic parentage dataset samples *Plectropomus maculatus,* a commercially- and recreationally-targeted species of grouper (Serranidae). The species is common in the approximately 700 ha of fringing reefs that are found in the Keppel Islands.

An earlier dataset (Harrison *et al.* 2012) reported adult and juvenile samples of *P. maculatus,* collected during 2007-2008, and these data have since been supplemented by additional sampling, and re-analysis of previously sampled juveniles with an enlarged adult dataset (Williamson *et al.*). The new dataset contains 440 adults and 506 juveniles from sampling efforts at 28 sites across the group (Figure 1); 58 of the juveniles were assigned to at least one adult parent. The summary parentage matrix, the habitat area of each location, and the inter-reef distances are given in the *Supplementary Information*. We apply our kernel estimator to identify the best-fit larval dispersal kernel from the set of thin- and fat-tailed generalised Gaussian functions; bootstrap resampling of sample locations produced 95% confidence intervals around this fit.

## Power analyses for simulated reef fish datasets

The collection and analysis of parentage data is both expensive and multi-dimensional: a patch must be visited, and adults and/or juveniles caught, genotyped (sometimes destructively), and then genetically matched. Total sampling effort must be shared between adults and juveniles, and across multiple patches. We perform a series of simulation-based power analyses to assess how these decisions affect the accuracy of the best-fit dispersal kernel. To do this, we (1) simulate dispersal and the resulting genetic parentage relationships, (2) create subsamples of juveniles and adults, and (3) analyse these simulated datasets using the fitting method described above.

Simulating dispersal: We begin by simulating a dispersal event for a single coral reef fish species across all 321 patch reefs in the Cairns Management Region of the Great Barrier Reef. A real seascape allows the power analyses to reflect the variation in natural inter-patch distances. The species’ characteristics are broadly based on coral trout (genus *Plectropomus*) species, a target of recreational fishers and a focus of genetic parentage analysis and larval kernel estimation (Harrison *et al.* 2012; Almany *et al.* 2013; Hopf *et al.* 2015). Each reef produces an amount of larvae proportional to its area, assuming that adult coral trout on the GBR exist at densities of around 3,500 km^-^2 (Cornish & Kiwi 2006), and mate randomly. We model the distance travelled by the larvae – which are obligate dispersers – using a random choice from a set of four variants of the generalised Gaussian (Eq. 1) for *θ* = {1, 2, 3, 0.5}. The chosen dispersal kernel is parameterised with a value of *k* that gives a mean dispersal distance *m* that is either “long” (*m* = 15 km) or “short” (*m* = 5km). The first three kernels are thin-tailed, the last is fat-tailed (see Figure S2 for both kernel shapes and parameter values). Not all larvae that arrive on a given reef survive post-recruitment density-dependent mortality, which we model using the Beverton-Holt function (Bode *et al.* 2012). We assume that mortality applies to settling larvae independent of their origin reef (but see: Burgess *et al.* 2013a).

This dispersal simulation creates a two-generation parentage dataset that can be repeatedly sampled in a simulation-based power analysis. A subset of the reefs within a contiguous region is randomly chosen for sampling (Figure 1A); from each reef we sample a proportion of the adult population and a given number of individuals from the post-density-dependent juvenile population. We use the average density of coral trout on the GBR to translate a proportional adult sample into a number of adult individuals (e.g., 1% of adults on a 1 km^2^ reef is equivalent to 35 individuals).

Fitting sample parentage data: We take each juvenile in turn and identify its source reef, if either parent was sampled. Although individual larvae will depart from and arrive at specific locations within each reef, we measure dispersal distances using the centroids of the source and destination reefs. While incorporating the precise locations of the adults and juveniles may hypothetically offer a more precise parameterisation, (1) inter-reef distances are generally much larger than reef dimensions, (2) dispersers are not necessarily spawned at the location where an adult was sampled (e.g., for aggregative spawners), and (3) juveniles did not necessarily settle at the precise location where they were sampled (White 2015). Parentage data are summarised in a simulated parentage matrix **M**, with the final row containing those juveniles whose parents were not in the sampled set.

We use our estimator (Eq. 4) to find the maximum likelihood parameter fit to these data, using the four candidate dispersal kernel functional forms. We repeat the sampling and fitting procedure 1,000 times to estimate confidence intervals, taking each sample from a different part of the metapopulation to average over the effects of a heterogeneous patch distribution (Figure 1A). We then calculate and report the mean dispersal distance *m* of the fitted kernel, and calculate how frequently the estimator identifies the true kernel shape (expressed as a percentage of the simulated datasets).

Power analyses: Our first power analysis assesses the performance of parentage analyses undertaken at the intensity and scale of contemporary sampling efforts. The case-studies in Table 1 involve sampling of between approximately 750 and 7,000 individuals (adults and juveniles) across 16 to 66 sites. We calculate the ability of this level of sampling to accurately estimate the two mean dispersal distances. We apply three scenarios of total sampling intensity – low sampling effort (750 individuals across 20 patches); intermediate effort (2,400 individuals across 40 patches); and high effort (6,000 individuals across 60 patches). In each case, total sampling effort is equally shared between adults and juveniles. At the same, we also calculate the performance of directly fitting a kernel to assigned juveniles (Telfer *et al.* 2003; Buston *et al.* 2012), ignoring all the factors we have discussed to this point. We simply calculate the maximum likelihood kernel fit, using the dispersal kernel to calculate the likelihood of each event (an assigned juvenile) as a function of the distance between parent(s) and assigned juvenile. We also estimate the accuracy of each case-study in Table 1, by calculating the width of the 95% confidence bounds, assuming that the best-fit parameter was correct.

**Table 1:** Details of case studies used to define parameter ranges. All values are taken from the references or their supplementary information. Juvenile sample column indicates number of samples, with the number and percentage of parentage assignments in parentheses. Adult column shows the number and estimated percentage of the adult population sampled. The final column shows the estimated accuracy of each dataset, measured with 95% confidence intervals, if the methods described in this paper were applied. *Each site in the Kimbe bay dataset contained multiple locations that other studies would have classified as sites. ** Except for one site, of which 50% were sampled. #We assumed that both juveniles and adults were sampled from the same set of patches. PNG = Papua New Guinea.

| Parentage analysis study | Sampling scale (approx.) | Adults |  | Juveniles |  | Best-fit kernel |  |  | 95% Confidence intervals |
| --- | --- | --- | --- | --- | --- | --- | --- | --- | --- |
| | | Total # | Sites | Total # | Sites | Kernel | $m$ | $\ln(k)$ | |
| *P. maculatus*<br>Keppels Island Group, Australia | 30 km | 466<br>(30%) | 19 | 493<br>(58; 12%) | 19 | Laplacian | 12.2 km | -2.5 | 5.9 – 71.3 km |
| *Elacatinus lori*<br>Carrie Bow Cay, Belize | 41 km | 3,033<br>(3%) | 30 | 4,122<br>(120; 3%) | 30 | Laplacian | 2.8 km | -1.02 | 2.7 – 2.9 km |
| *P. areolatus*<br>Manus Island, PNG | 55 km | 416<br>(43%) | 1 | 782<br>(76; 10%) | 66 | Ribbens | 14.5 km | -3.4# | 6.4 – 26.4 km |
| *Amphiprion percula*<br>Kimbe Bay, PNG | 50 km | 506<br>(100%) | 1 | 469<br>(122; 26%) | 4* | N/A | N/A | N/A | N/A |
| *Stegastes partitus*<br>Exuma Sound, Bahamas | 140 km | 314<br>(N/A) | 11 | 437<br>(2; <1%) | 7 | N/A | N/A | N/A | N/A |
| *A. polymnus*<br>Bootless bay, PNG | 30 km | 452<br>(100%**) | 8 | 491<br>(100; 20%) | 8 | N/A | N/A | N/A | N/A |

The remaining power analyses aim to inform how a given sampling budget should best be distributed. In general, total parentage sampling effort is distributed along three primary dimensions: the number of sampled (1) juveniles, (2) adults, and (3) sites. Total sampling effort is the product of these three; different effort allocations affect the accuracy of the estimated kernel. We outline four additional power analyses below, and on the basis of the results recommend how total sampling effort should be distributed to best estimate the kernel shape and the mean dispersal distance. For all power analyses we calculate the performance of a range of total sampling intensities, comprising 500, 1000, 2500, 3250 and 5000 individuals, and consider species with both long- and short-distance dispersal kernels.

Power analysis 2 considers how to best distribute sampling effort between juveniles and adults. A focus on sampling juveniles generates a larger number of dispersal events, however, more adult samples give a higher probability that each event will be usefully attributed to a source and destination reef. Assuming that both adults and juveniles are sampled from the same set of 30 reefs, we sample the genetics individuals spread equally across reefs. We share this sampling effort between adults and juvenile individuals in different proportions, ranging from 5% adults (and therefore 95% juveniles) through to 95% adults.

In power analysis 3, we consider the distribution of sampling effort across space. Sampling across a larger number of patches increases our ability to sample the kernel tail (D’Aloia *et al.* 2015), but it means that we sample fewer individuals on each patch, and therefore see more unassigned juveniles. Assuming that sampling will be equally focused (50:50) on adults and juveniles, we sample the genetics of individuals collected from between 5 and 90 different patches. We note that, while the spatial scale of the sampling will scale on average with the number of patches (i.e., a larger number of patches is generally distributed over a larger area), the absolute scale in kilometers will depend on the density of patches in the sample area (Figure 1A).

In power analysis 4, we consider the implications of incomplete sampling of adult populations on sampled patches. When a sampled juvenile cannot be assigned to one or both parents, the kernel fitting procedure must decide if it came from a ghost patch (usually a long-distance disperser), or from unsampled adults on a sampled patch (usually a short-distance disperser or self-recruiter). Low intensity adult sampling sends conflicting messages to the estimator because it makes both options a possibility. We sample 500 juveniles across 30 reefs, and apply adult sampling proportions that range between 1% and 100%. Note that, unlike the previous power analyses, these simulations have unequal total sampling numbers. The fit will therefore improve monotonically as adult sampling increases.

Power analysis 5 calculates the impact of ignoring ghost patches, unsampled adults and unassigned juveniles into our fitting methods. We create parentage data for sampled individuals distributed equally across adults and juveniles on 30 reefs. We contrast the mean dispersal distance estimated using the methods we propose in Eq. 4, or estimated by fitting a dispersal kernel to the assigned samples only, without considering unsampled factors. That is, we ignore (1) the unassigned juveniles, (2) the unsampled patches, and (3) the presence of unsampled adults on sampled patches.

## Results

Figure 1C illustrates the best-fit kernels for *P. maculatus* dispersal, measured by the Keppel Islands dataset. The data was best fit with the Ribbens kernel (Eq. 1 with *θ* = 3), but each of the thin-tailed kernels predicts an approximately equal mean dispersal distance (22.5 km, 24.2 km, 26.9 km and 35.1 km for *θ* = 1, 2, 3 and 4 respectively). The best-fit Gaussian kernel (*θ* = 2) fit within the 95% confidence intervals of the best-fit Ribbens kernel, and had a relatively strong AIC weight (0.15). Each of these best-fit kernels predicts that the majority of larval settlement within the Keppels group is sourced from other reefs in the group, and also that a large proportion of the larval output of the group recruits to local reefs. The final column in Table 1 indicates the estimated accuracy of three different datasets that have been used to fit larval dispersal kernels. Only the parentage dataset of *Elacatinus lori* is sufficiently powerful to produce narrow confidence bounds around the mean dispersal distance of the species, if the methods we outline here were applied.

In our first power analysis, we calculate the expected performance of current levels of sampling (Figure 2). For a species with a mean dispersal distance of 15 km, low sampling effort (1,000 individuals across 20 patches) provides an estimate between 5.7 km and 42.6 km (38% to 285% of the true value); intermediate effort (2,000 individuals across 40 patches) provides an estimate between 9.5 km and 28.7 km (64% to 191%); and high effort (7,000 individuals across 70 patches) provides an estimate between 13.2 km and 18.9 km (88% to 126%). The results are more precise for the species with shorter mean dispersal distance of 5 km, where low sampling effort provides an estimate between 2.5 km and 12.0 km (50% to 240%); intermediate effort provides an estimate between 4.2 km and 6.4 km (83% to 130%); and high effort provides an estimate between 4.7 km and 5.5 km (94% to 111%). In all cases, the straightforward approach of directly fitting kernels to observed parentage assignments provided poor, biased underestimates of mean dispersal distance (Figure 2). The 95% confidence intervals only once included the true distance (for high-intensity sampling of long distance dispersers).

**Figure 2:**
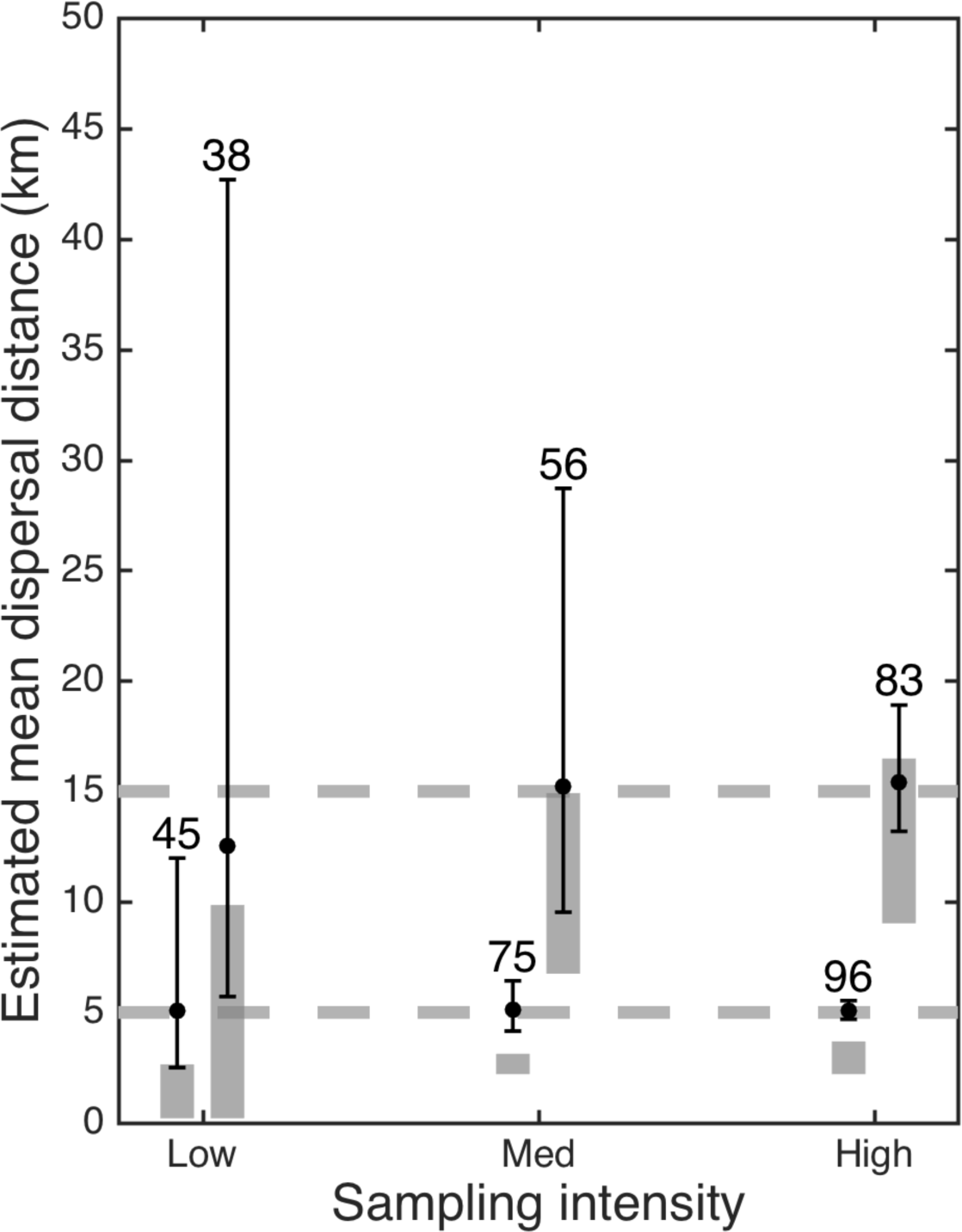
Accuracy of the kernel estimator for three different sampling intensities: low (750 individuals across 20 patches); intermediate (2,400 individuals across 40 patches); and high (6,000 individuals across 60 patches). Error bars enclose 95% of the power analysis simulations. Shaded areas indicate the results of fitting dispersal kernels directly to the distances travelled by assigned juveniles (areas enclose 95% of the simulations). Left hand results for each sampling intensity are fitting the short-distance (5 km) disperser; right hand bars are fitting the long-distance (15 km) disperser.

Our second power analysis considers the distribution of sampling effort between juveniles and adults (Figure 3 A-B). The performance of the estimator does not vary substantially with this proportion allocation. The best results are achieved by allocating the sampling effort evenly, but the outcomes are essentially the same, as long as allocations do not fall below 25% of sampling effort for either adults or juveniles. Datasets outside this range tend to provide biased estimates of the mean dispersal distance, particularly for longer distance dispersers.

**Figure 3:**
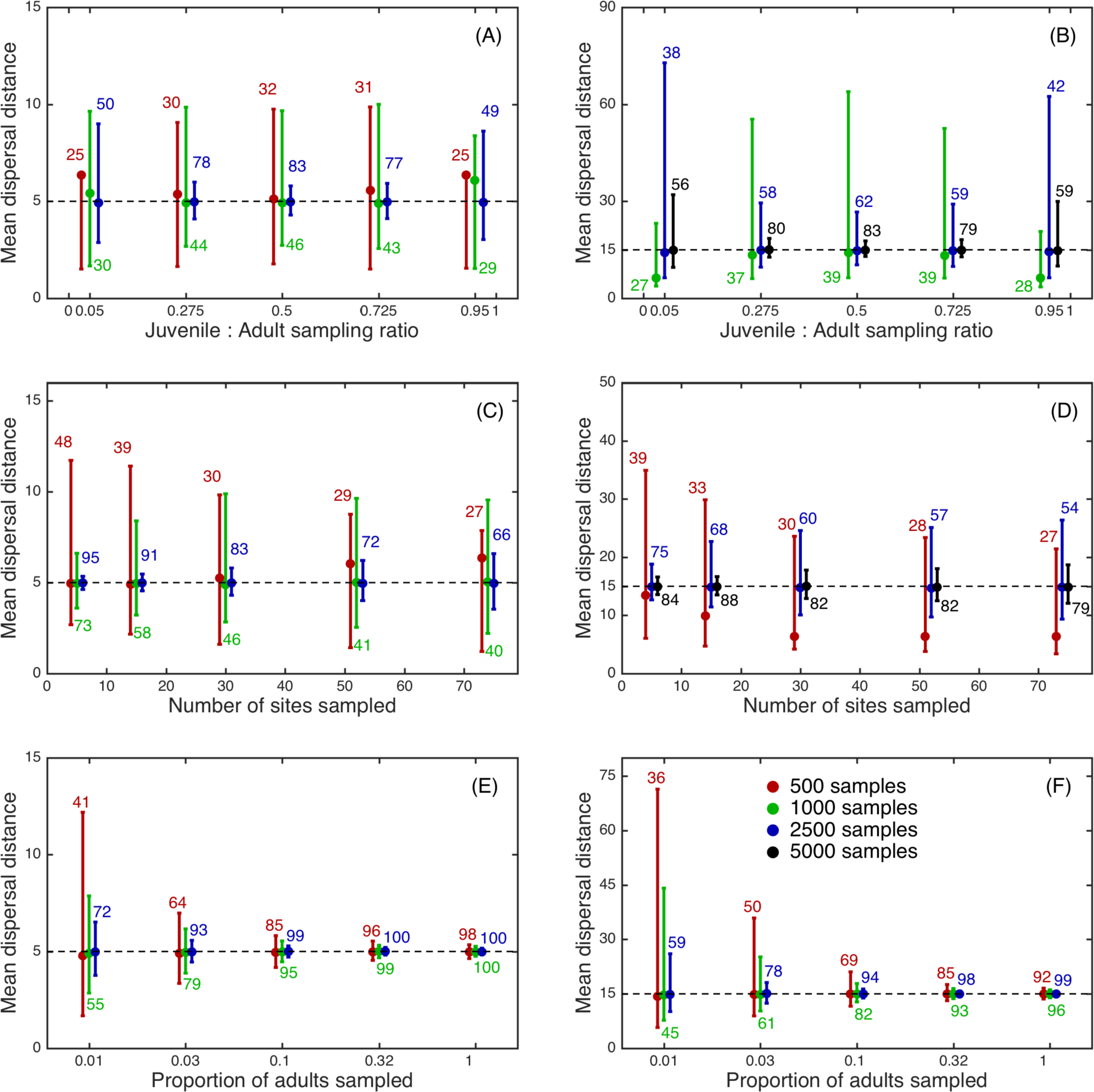
Accuracy of the kernel estimator for different allocations of sampling effort, measuring mean dispersal distance (y-axis). Different panels show the effect of distributing total sampling effort in different ways between adults and juveniles (A-B), and among a different number of reefs (C-D); and the effects of sampling larger proportions of the adult population (E-F). Colours show different total sampling intensities (legend in panel F, for clarity, not all are shown in each panel). Panels in the left column (A, C,E) are for the short-distance (5 km) disperser; the right column (B, D,F) shows the long-distance (15 km) disperser. Dashed line indicates the true mean dispersal distance. All error bars enclose 95% of the simulations. Numbers indicate the percentage of simulations where the true kernel shape was correctly identified.

The third analysis considers the distribution of sampling effort across space. The most accurate and precise estimates of dispersal distances are achieved by focusing sampling effort on a small number of well-sampled patches. Sampling individuals over a wider area results in a less precise estimate of the kernel, for both long- and short-distance dispersers (Figure 3 C-D). Spreading a small number of samples over a large space results in substantial underestimates of the mean dispersal distance. For larger sampling budgets, the size of the sampling region has a smaller influence on the fit precision.

The fourth analysis addresses incomplete adult sampling. The results show that the estimator is unbiased, regardless of the proportion of the adult population sampled (Figure 3 E-F), and is consistently precise if the proportion of the adult population sampled remains above 5%. However, once the proportion fell below this level, the performance of the fit declined dramatically, particularly for lower total sampling budgets. Unlike the previous figures, this power analysis keeps all other sampling decisions constant as it increases the proportion of adults sampled. These results would therefore not justify increasing the sampled adult proportion above 10%, since the additional adults would have little impact on the fit, but would reduce the number of juveniles or patches sampled.

The final power analysis measures the impact of ignoring ghost patches and unassigned juveniles when fitting kernels. Not including these populations has a negative effect on estimator performance – specifically, it causes large underestimates – particularly when fewer individuals were sampled, and for short dispersers (Figure 4). In conditions when the inclusion of unsampled patches produced accurate and precise estimates of the dispersal kernel (i.e., 5,000 individuals sampled across 30 reefs), ignoring the effects of unsampled adults meant that the 95% confidence intervals no longer enclosed the true value.

**Figure 4:**
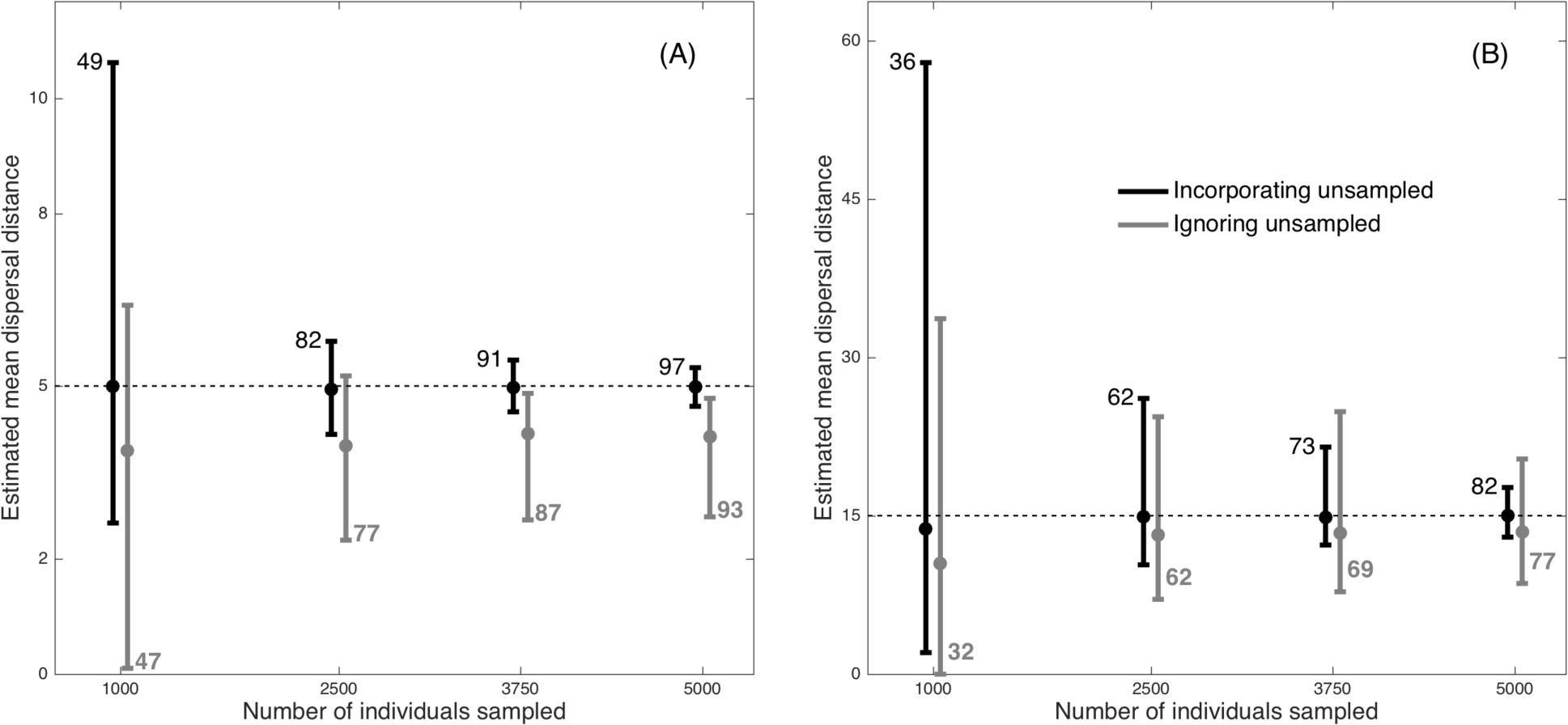
Effect of incorporating partial sampling (unsampled adults, ghost populations, and unassigned juveniles) into the kernel estimator. Black lines show results for the estimator that includes partial sampling; grey lines show results when the estimator ignores partial sampling. Numbers denote the percentage of simulations where the kernel shape was correctly identified. Dashed line shows the true mean dispersal distance for the short-distance (A) and long-distance (B) disperser; error bars enclose 95% simulated parentage datasets. Results are shown for four levels of total sampling effort, distributed evenly between adults and juveniles across 30 reefs.

## Discussion

Our power analyses demonstrate that, when fitting dispersal kernels and estimating dispersal distance, it is essential to account for the presence of ghost patches and unsampled adults. Fitted relationships between distance and dispersal strength that do not consider these adults, and particularly direct fits of kernels to assigned juveniles, are likely to be inaccurate. Our results indicate that to accurately characterise a dispersal kernel using parentage analyses, researchers should ideally collect at least 5,000 samples from around 10 patches. The samples should be allocated as evenly as possible between adults and juveniles, while ensuring that at least 5% of the adults on each patch are sampled. While quantitatively useful for planning future estimates, many of our conclusions are qualitatively unsurprising. For example, greater total sampling effort produces better quality fits, as does sampling larger proportions of the adult population on each reef. Other conclusions were less immediately obvious. For example, we show that total effort is best distributed relatively equally between adults and juveniles. However, a strong emphasis on adults would result in a higher proportion of assignments, which could plausibly have given better estimates of the kernel. Sampling is also best concentrated on a relatively small number of sites compared to the dispersal ability of the species, greatly increasing the likelihood of positive assignments. However, it was also intuitively possible that an intermediate number of patches would have performed best, since this would have given a broader cross-section of the kernel shape, including the tail.

On the basis of these results, only some of the currently available parentage datasets are powerful enough to provide reasonably accurate estimates of species’ dispersal kernels (Table 1). While some studies have sampled in excess of 10% of the adults (Saenz-Agudelo *et al.* 2011; Almany *et al.* 2013), others are much lower or unknown (Christie *et al.* 2010; D’Aloia *et al.* 2015). Many have achieved sample sizes in excess of 5000 (D’Aloia *et al.* 2015), but others have been based in very limited sampling (Christie *et al.* 2010; Harrison *et al.* 2012). While our results show that current datasets are large enough to accurately characterise dispersal kernels, this is only true if managers apply appropriate fitting methods. In particular, we focus on two improvements to previous approaches: first, we incorporate unassigned juveniles, both by including the influence of ghost patches, and by correcting for unsampled adults on partially sampled reefs. Although these factors are understood to be important (Beerli 2004; Jones *et al.* 2005a; Wang 2014), they have not yet been incorporated into kernel estimators for species in coral reef ecosystems, where both are ubiquitous. Second, we acknowledge that the juveniles in parentage datasets are sampled after post-settlement density dependence has occurred. As a result, the proportion of dispersers who travel from patch *i* to patch *j* (dispersal: the predictions of a kernel) is very different the proportion of juveniles on patch *j* who can be assigned to adults on patch *i* (recruitment: the observations of a parentage analysis). This difference between dispersal and recruitment was highlighted by Burgess *et al.* (2014); our results (e.g., Supplementary Figure S1) demonstrate that the difference must also be taken into account when fitting dispersal kernels.

Although our method includes more factors than previous efforts, it nevertheless makes simplifying assumptions about the processes that generate parentage data. First, we assume that parentage assignment is completely accurate, but the allocation error of parent-offspring relationships for reef fish species has been estimated at <5% for false-positives, and <1% for false-negatives (Harrison *et al.* 2012). While very low, these mis-assignments will lower the precision of our estimates. Moreover, the amount of sampling effort directed at the adult populations will affect these rates (Harrison *et al.* 2013b; Christie 2013; see: Harrison *et al.* 2013a) independent of our modelled effects of sample size on precision. These facts should be incorporated into the fit statistic. The likely result will be an increase in the uncertainty of the estimates but not their bias, and a greater emphasis on sampling parents than is current seen in our results. Second, the estimator assumes binomial sampling with replacement, implying that the settlement pool is much larger than the number of settlers, and that the number of juvenile samples is a small proportion of the total number of juveniles. Larger proportions would require a different sampling model, and some estimate of the total number of settlers and juveniles. Third, we assume a great deal of spatial homogeneity in a series of key dispersal processes: mortality in the pelagic environment and upon settlement; adult density and fecundity; and the strength of density dependence. Such assumptions are probably incorrect in most ecosystems, although evidence is generally scarce. For reef fish species, each of these processes is thought to vary in space, and/or with the origin of the dispersers (Ruttenberg *et al.* 2005; Sponaugle & Grorud-Colvert 2006; Burgess *et al.* 2013b; Hixon & Webster), although the amount and patterns of variation are unknown for almost all species in almost all locations. Our assumptions of homogeneity therefore reflect data limitations. Finally, our estimator only predicts proportional dispersal kernels, rather than the absolute number of spawned individuals who travel a given distance. The latter dispersal kernels cannot be constructed without additional information – specifically, estimates of settlement numbers before density-dependent mortality occurs – a very difficult quantity to measure (Almany & Webster 2006).

Each of these assumptions and simplifications could be corrected with alterations to the estimator, if appropriate data could be collected. However, a much broader concern is the applicability and utility of kernel descriptions of dispersal in ecology. This is a particular concern for coral reef fish, whose ecology is governed by obligate dispersal, and which are highly dispersive species in an advective and turbulent environment. Kernels use smooth probability distributions, which essentially assume that the dispersal process is homogeneous and temporally-consistent. Propagules radiate out from each natal patch in a spatially-invariant and isotropic pattern; where advection is included, it is modelled as a consistent displacement of this smooth pattern (Almany *et al.* 2013; D’Aloia *et al.* 2015). The strength of dispersal to nearby patches is determined entirely by the distance between them, but in reality the density of habitat patches will influence dispersal, as locomotive individuals choose between different settlement options (Gerlach *et al.* 2007), or choose to delay settlement in the hope of finding more suitable habitat. Importantly, biophysical models of larval dispersal, based on a complex understanding of oceanographic processes and forced and validated by extensive data, predict dispersal patterns that are highly variable in both space and time, at multiple scales (James *et al.* 2002; Cowen 2006; Bode *et al.* 2012). It is possible that biophysical dispersal – if averaged over a sufficiently long timespan – approaches a smooth kernel (Cowen & Sponaugle 2009), and that long-term management decisions can be based on such time-averaged descriptions of dispersal. However, it is currently unclear whether coral reef fish dispersal in particular, and ecological dispersal in general, can be accurately described using dispersal kernels. On the other hand, the inaccuracy of kernel descriptions of dispersal may not automatically preclude them being useful for management. If smooth kernels capture essential elements of dispersal dynamics – for example, the mean distance travelled – then decisions based on these imperfect descriptions may still satisfy management objectives if they respond primarily to mean dispersal distance (Runge *et al.* 2011; Moore & Runge 2012). That is, kernels may not accurately capture all the processes of dispersal, but they may reflect enough of the underlying dynamics to plan marine reserve networks (HALPERN *et al.* 2006), or manage fisheries (Bode *et al.* 2016).

The contrast between the biophysical and kernel descriptions of dispersal therefore creates an important challenge for spatial ecologists and conservation biologists. Mechanistic, biophysical models of dispersal are almost certainly more accurate than the kernel-based descriptions, and explicitly include factors (e.g., spatial variation; temporal fluctuations) that are known to fundamentally alter ecological processes and conservation management (Chesson & Warner 1981; Amarasekare 2003; Berkeley *et al.* 2010). However, most ecological (Bode *et al.* 2011; Okubo & Levin 2013), evolutionary (Skellam 1951; Levin *et al.* 2003; Nathan 2006) and management theory (Hastings & Botsford 2003; Neubert & Parker 2004; Arim *et al.* 2006; White *et al.* 2008; Botsford *et al.* 2009), and almost all of our spatial planning tools (Moilanen & Wintle 2006; Lehtomaki *et al.* 2009; Carroll *et al.* 2010; Laitila & Moilanen 2013), are based on dispersal kernels, rather than connectivity matrices (the output of biophysical models). These two descriptions of dispersal must be reconciled, or the simpler, kernel-based description must be refuted. This is especially true for the understanding and management of reef fish biodiversity, where dispersal plays such a pivotal role in demography (Cowen 2002; Bode *et al.* 2006), community dynamics (Salomon *et al.* 2010; Bode *et al.* 2011), and spatial management (Hastings & Botsford 2003; Almany *et al.* 2013; Green *et al.* 2014). Our development and testing of accurate dispersal kernel fitting methods is an essential step in this process.

## References

Almany, G.R. & Webster, M.S. (2006). The predation gauntlet: early post-settlement mortality in reef fishes. Coral Reefs, 25, 19–22.

Almany, G.R., Berumen, M.L., Thorrold, S.R., Planes, S. & Jones, G.P. (2007). Local Replenishment of Coral Reef Fish Populations in a Marine Reserve. Science, 316, 742–744.

Almany, G.R., Hamilton, R.J., Bode, M., Matawai, M., Potuku, T., Saenz-Agudelo, P., Planes, S., Berumen, M.L., Rhodes, K.L., Thorrold, S.R., Russ, G.R. & Jones, G.P. (2013). Dispersal of Grouper Larvae Drives Local Resource Sharing in a Coral Reef Fishery. Current Biology, 23, 626–630.

Amarasekare, P. (2003). Competitive coexistence in spatially structured environments: a synthesis. Ecology Letters, 6, 1109–1122.

Arim, M., Abades, S.R., Neill, P.E., Lima, M. & Marquet, P.A. (2006). Spread dynamics of invasive species. Proceedings of the National Academy of Sciences of the United States of America, 103, 374–378.

Beerli, P. (2004). Effect of unsampled populations on the estimation of population sizes and migration rates between sampled populations. Molecular Ecology, 13, 827–836.

Berkeley, H.A., Kendall, B.E., Mitarai, S. & Siegel, D. (2010). Turbulent dispersal promotes species coexistence. Ecology Letters, 13, 360–371.

Bode, M., Armsworth, P.R., Fox, H.E. & Bode, L. (2012). Surrogates for reef fish connectivity when designing marine protected area networks. Marine Ecology Progress Series, 466, 155–166.

Bode, M., Bode, L. & Armsworth, P.R. (2011). Different dispersal abilities allow reef fish to coexist. Proceedings of the National Academy of Sciences, 108, 16317–16321.

Bode, M., Bode, L. & Armsworth, P.R. (2006). Larval dispersal reveals regional sources and sinks in the Great Barrier Reef. Marine Ecology Progress Series, 308, 17–25.

Bode, M., Sanchirico, J.N. & Armsworth, P.R. (2016). Returns from matching management resolution to ecological variation in a coral reef fishery. Proceedings of the Royal Society B: Biological Sciences, 283, 20152828.

Botsford, L.W., White, J.W., Coffroth, M.A., Paris, C.B., Planes, S., Shearer, T.L., Thorrold, S.R. & Jones, G.P. (2009). Connectivity and resilience of coral reef metapopulations in marine protected areas: matching empirical efforts to predictive needs. Coral Reefs, 28, 327–337.

Broquet, T. & Petit, E.J. (2009). Molecular Estimation of Dispersal for Ecology and Population Genetics. dx.doi.org.ezp.lib.unimelb.edu.au, 40, 193–216.

Bullock, J.M., Kenward, R.E. & Hails, R.S. (2002). Dispersal ecology: 42nd symposium of the British ecological society.

Burgess, S.C., Bode, M. & Marshall, D.J. (2013a). Costs of dispersal alter optimal offspring size in patchy habitats: combining theory and data for a marine invertebrate (C. Fox, Ed.). Functional Ecology, 27, 757–765.

Burgess, S.C., Bode, M. & Marshall, D.J. (2013b). Costs of dispersal alter optimal offspring size in patchy habitats: combining theory and data for a marine invertebrate (C. Fox, Ed.). Functional Ecology, 27, 757–765.

Burgess, S.C., Nickols, K.J., Griesemer, C.D., Barnett, L.A.K., Dedrick, A.G., Satterthwaite, E.V., Yamane, L., Morgan, S.G., White, J.W. & Botsford, L.W. (2014). Beyond connectivity: how empirical methods can quantify population persistence to improve marine protected-area design. Ecological Applications, 24, 257–270.

Buston, P.M., Jones, G.P., Planes, S. & Thorrold, S.R. (2012). Probability of successful larval dispersal declines fivefold over 1 km in a coral reef fish. Proceedings of the Royal Society B: Biological Sciences, 279, 1883–1888.

Carroll, C., Dunk, J.R. & Moilanen, A. (2010).Optimizing resiliency of reserve networks to climate change: multispecies conservation planning in the Pacific Northwest, USA. Global Change Biology, 16, 891–904.

Chesson, P.L. & Warner, R.R. (1981). Environmental variability promotes coexistence in lottery competitive systems. The American Naturalist, 117, 923–943.

Christie, M.R. (2013). Bayesian parentage analysis reliably controls the number of false assignments in natural populations. Molecular Ecology, 22, 5731–5737.

Christie, M.R., Johnson, D.W., Stallings, C.D. & Hixon, M.A. (2010). Self-recruitment and sweepstakes reproduction amid extensive gene flow in a coral-reef fish. Molecular Ecology, 19, 1042–1057.

Clark, J.S., Silman, M., Kern, R., Macklin, E. & HilleRisLambers, J. (2008). SEED DISPERSAL NEAR and FAR: PATTERNS ACROSS TEMPERATE and TROPICAL FORESTS. dx.doi.org.ezp.lib.unimelb.edu.au, 80, 1475–1494.

Clobert, J., Danchin, E., Dhondt, A.A. & Nichols, J.D. (2001). Dispersal.

Cornish, A. & Kiwi, L.K. (2006). Plecropomus leopardus. Species Report for the IUCN Red List of Threatened Species. IUCN, Gland, Switzerland.

Cowen, R.K. (2002). Larval dispersal and retention and consequences for population connectivity. Coral reef fishes: dynamics and diversity in a complex ecosystem (ed P. Sale), pp. 149–170. Academic Press, San Diego.

Cowen, R.K. (2006). Scaling of Connectivity in Marine Populations. Science, 311, 522–527.

Cowen, R.K. & Sponaugle, S. (2009). Larval dispersal and marine population connectivity. Annual Review of Marine Science, 1, 443–466.

D’Aloia, C.C., Bogdanowicz, S.M., Francis, R.K., Majoris, J.E., Harrison, R.G. & Buston, P.M. (2015). Patterns, causes, and consequences of marine larval dispersal. Proceedings of the National Academy of Sciences, 112, 13940–13945.

Ellstrand, N.C. & Marshall, D.L. (1985). Interpopulation gene flow by pollen in wild radish, Raphanus sativus. American Naturalist.

Gerlach, G., Atema, J., Kingsford, M.J., Black, K.P. & Miller-Sims, V. (2007). Smelling home can prevent dispersal of reef fish larvae. Proceedings of the National Academy of Sciences of the United States of America, 104, 858–863.

Green, A.L., Fernandes, L. & Almany, G. (2014). Designing marine reserves for fisheries management, biodiversity conservation, and climate change adaptation. Coastal ….

Halpern, B.S., Regan, H.M., Possingham, H.P. & McCarthy, M.A. (2006). Accounting for uncertainty in marine reserve design. Ecology Letters, 9, 2–11.

Harrison, H.B., Saenz-Agudelo, P., Planes, S., Jones, G.P. & Berumen, M.L. (2013a). On minimizing assignment errors and the trade-off between false positives and negatives in parentage analysis. Molecular Ecology, 22, 5738–5742.

Harrison, H.B., Saenz-Agudelo, P., Planes, S., Jones, G.P. & Berumen, M.L. (2013b). Relative accuracy of three common methods of parentage analysis in natural populations. Molecular Ecology, 22, 1158–1170.

Harrison, H.B., Williamson, D.H., Evans, R.D., Almany, G.R., Thorrold, S.R., Russ, G.R., Feldheim, K.A., van Herwerden, L., Planes, S., Srinivasan, M., Berumen, M.L. & Jones, G.P. (2012). Larval Export from Marine Reserves and the Recruitment Benefit for Fish and Fisheries. Current Biology, 22, 1023–1028.

Hastings, A. & Botsford, L.W. (2003). Comparing designs of marine reserves for fisheries and for biodiversity. Ecological Applications, 13, S65–S70.

Hixon, M. & Webster, M.S. Density dependence in reef fish populations. Coral Reef Fishes: Dynamics and Diversity in a Complex Ecosystem (ed P.F. Sale), pp. 303–325. Academic Press, San Diego, California.

Hopf, J.K., Jones, G.P., Willamson, D.H. & Connolly, S.R. (2015). Fishery consequences of marine reserves: short-term pain for longer-term gain. Ecological Applications, 26, 818–829.

James, M.K., Armsworth, P.R., Mason, L.B. & Bode, L. (2002). The structure of reef fish metapopulations: modelling larval dispersal and retention patterns. Proceedings of the Royal Society B: Biological Sciences, 269, 2079–2086.

Jones, A.G. & Ardren, W.R. (2003). Methods of parentage analysis in natural populations. Molecular Ecology, 12, 2511–2523.

Jones, F.A. & Muller Landau, H.C. (2008). Measuring long-distance seed dispersal in complex natural environments: an evaluation and integration of classical and genetic methods. Journal of Ecology, 96, 642–652.

Jones, F.A., Chen, J., Weng, G.J. & Hubbell, S.P. (2005a). A Genetic Evaluation of Seed Dispersal in the Neotropical Tree Jacaranda copaia(Bignoniaceae). The American Naturalist, 166, 543–555.

Jones, G.P., Almany, G.R., Russ, G.R., Sale, P.F., Steneck, R.S., Oppen, M.J.H. & Willis, B.L. (2009). Larval retention and connectivity among populations of corals and reef fishes: history, advances and challenges. Coral Reefs, 28, 307–325.

Jones, G.P., Planes, S. & Thorrold, S.R. (2005b). Coral Reef Fish Larvae Settle Close to Home. Current Biology, 15, 1314–1318.

Laitila, J. & Moilanen, A. (2013). Approximating the dispersal of multi-species ecological entities such as communities, ecosystems or habitat types. Ecological Modelling, 259, 24–29.

Largier, J.L. (2003). Considerations in estimating larval dispersal distances from oceanographic data. Ecological Applications.

Lehtomaki, J., Tomppo, E., Kuokkanen, P., Hanski, I. & Moilanen, A. (2009). Applying spatial conservation prioritization software and high-resolution GIS data to a national-scale study in forest conservation. Forest Ecology and Management, 258, 2439–2449.

Levin, S.A., Muller-Landau, H.C. & Nathan, R. (2003). The ecology and evolution of seed dispersal: a theoretical perspective. Annual Review of Ecology.

Moilanen, A. & Wintle, B.A. (2006). Uncertainty analysis favours selection of spatially aggregated reserve networks. Biological Conservation, 129, 427–434.

Moore, J.L. & Runge, M.C. (2012). Combining Structured Decision Making and Value-of-Information Analyses to Identify Robust Management Strategies. Conservation Biology, 26, 810–820.

Moran, E.V. & Clark, J.S. (2012). Between-Site Differences in the Scale of Dispersal and Gene Flow in Red Oak (B. Fenton, Ed.). PLoS ONE, 7, e36492.

Nathan, R. (2006). Long-Distance Dispersal of Plants. Science, 313, 786–788.

Nathan, R., Klein, E., Robledo-Arnuncio, J.J. & Revilla, E. (2012). Dispersal kernels: review. Dispersal Ecology and Evolution (eds J. Clobert, M. Baguette, T.G. Benton & J.M. Bullock), p. 496. Oxford University Press, Oxford.

Neubert, M.G. & Parker, I.M. (2004). Projecting Rates of Spread for Invasive Species. Risk Analysis, 24, 817–831.

Okubo, A. & Levin, S.A. (2013). Diffusion and ecological problems: modern perspectives.

Planes, S., Jones, G. & Thorrold, S. (2009). Larval dispersal connects fish populations in a network of marine protected areas. Proceedings of the National Academy of Sciences, 106, 5693–5700.

Runge, M.C., Converse, S.J. & Lyons, J.E. (2011). Which uncertainty? Using expert elicitation and expected value of information to design an adaptive program. Biological Conservation, 144, 1214–1223.

Ruttenberg, B.I., Haupt, A.J., Chiriboga, A.I. & Warner, R.R. (2005). Patterns, causes and consequences of regional variation in the ecology and life history of a reef fish. Oecologia, 145, 394–403.

Saenz-Agudelo, P., Jones, G.P., Thorrold, S.R. & Planes, S. (2011). Connectivity dominates larval replenishment in a coastal reef fish metapopulation. Proceedings of the Royal Society B: Biological Sciences, 278, 2954–2961.

Salomon, Y., Connolly, S.R. & Bode, L. (2010). Effects of asymmetric dispersal on the coexistence of competing species. Ecology Letters, 13, 432–441.

Skellam, J.G. (1951). Random Dispersal in Theoretical Populations. Biometrika, 38, 196.

Sponaugle, S. & Grorud-Colvert, K. (2006). Temperature-mediated variation in early life history traits and recruitment success of the coral reef fish Thalassoma bifasciatum in the Florida Keys. Marine Ecology Progress Series.

Telfer, S., Piertney, S.B., Dallas, J.F., Stewart, W.A., Marshall, F., Gow, J.L. & Lambin, X. (2003). Parentage assignment detects frequent and large-scale dispersal in water voles. Molecular Ecology, 12, 1939–1949.

Wang, J. (2014). Estimation of migration rates from marker-based parentage analysis. Molecular Ecology.

Warner, R.R. (1997). Evolutionary ecology: how to reconcile pelagic dispersal with local adaptation. Coral Reefs, 16, S115–S120.

White, C., Kendall, B.E., Gaines, S., Siegel, D.A. & Costello, C. (2008). Marine reserve effects on fishery profit. Ecol Lett, 11, 370–379.

White, J.W. (2015). Marine reserve design theory for species with ontogenetic migration. Biology Letters, 11, 20140511–20140511.

Williamson, D.H., Harrison, H.B., Almany, G.R., Berumen, M.L., Bode, M., Bonin, M.C., Choukroun, S., Doherty, P.J., Frisch, A.J., Saenz-Agudelo, P. & Jones, G.P. Large-scale, multi-directional larval connectivity among coral reef fish populations in the Great Barrier Reef Marine Park.

Woltmann, S., Sherry, T.W. & Kreiser, B.R. (2012). A genetic approach to estimating natal dispersal distances and self-recruitment in resident rainforest birds. Journal of avian biology, 43, 33–42.

